# Reducing voltage-dependent potassium channel Kv3.4 levels ameliorates synapse loss in a mouse model of Alzheimer’s disease

**DOI:** 10.1101/2021.11.24.469829

**Authors:** Jie Yeap, Chaitra Sathyaprakash, Jamie Toombs, Jane Tulloch, Cristina Scutariu, Jamie Rose, Karen Burr, Caitlin Davies, Marti Colom-Cadena, Siddharthan Chandran, Charles H Large, Matthew JM Rowan, Martin J Gunthorpe, Tara L Spires-Jones

**Affiliations:** UK Dementia Research Institute and Centre for Discovery Brain Sciences, University of Edinburgh, UK; UK Dementia Research Institute and Centre for Clinical Brain Sciences, University of Edinburgh UK; Autifony Therapeutics Limited, Stevenage Bioscience Catalyst, Stevenage, UK; Emory University School of Medicine, Atlanta, GA, USA

## Abstract

Synapse loss is associated with cognitive decline in Alzheimer’s disease (AD) and owing to their plastic nature, synapses are an ideal target for therapeutic intervention. Oligomeric amyloid beta (Aβ) around amyloid plaques is known to contribute to synapse loss in mouse models and is associated with synapse loss in human AD brain tissue, but the mechanisms leading from Aβ to synapse loss remain unclear. Recent data suggest that the fast-activating and -inactivating voltagegated potassium channel subtype 3.4 (K_v_3.4) may play a role in Aβ-mediated neurotoxicity. Here, we tested whether this channel could also be involved in Aβ synaptotoxicity. Using adeno-associated virus and CRISPR (clustered regularly interspaced short palindromic repeats) technology, we reduced K_v_3.4 expression in neurons of the somatosensory cortex of APP/PS1 mice. These mice express human familial AD associated mutations in amyloid precursor protein and presenilin 1 and develop amyloid plaques and plaque-associated synapse loss similar to that observed in AD brain. We observe that reducing K_v_3.4 levels ameliorates dendritic spine loss and changes spine morphology compared to control virus. In support of translational relevance, K_v_3.4 protein was observed in human AD and control brain and is associated with synapses in human iPSC-derived cortical neurons. Interestingly, we observe a decrease in K_v_3.4 expression in iPSC derived cortical neurons when they are challenged with human Alzheimer’s disease derived brain homogenate. These results suggest that approaches to reduce K_v_3.4 expression and/or function could be protective against Aβ-induced synaptic alterations.

## Introduction

Alzheimer’s disease (AD) is characterised by the progressive accumulation of amyloid plaques and neurofibrillary tangles, composed of abnormally aggregated β-amyloid (Aβ) and hyperphosphorylated tau proteins, respectively (Serrano-Pozo et al., 2011). Along with these neuropathological lesions there is extensive neuronal loss, synapse loss, and gliosis (Henstridge et al., 2019). Of these brain changes, synapse loss is the strongest correlate of cognitive decline in AD (DeKosky & Scheff, 1990; Terry et al, 1991), and synapses, with their myriad receptors and ion channels, are attractive therapeutic targets for intervention (Colom-Cadena et al., 2020). In brains affected by AD and in mouse models of amyloid plaque formation, plaques act as a local reservoir of oligomeric Aβ, emanating these soluble and diffusible Aβ species to form a halo surrounding plaques which is associated with synapse loss (Koffie et al., 2009, 2012; Spires-Jones & Hyman, 2014). While the synaptotoxicity of oligomeric Aβ is well-established, it is not yet entirely clear how synapses are damaged or whether this can be recovered by therapeutic interventions.

One potential pathway mediating Aβ synaptotoxicity involves voltage-gated potassium channels. The voltage-gated potassium channel subtype 3.4 (K_v_3.4) encoded by the *KCNC4* gene is a member of the K_v_3 family of channels that play important roles in controlling neuronal firing and contribute to synaptic plasticity (Kaczmarek & Zhang, 2017; Rowan & Christie, 2017). This channel is believed to have vital roles in regulating neurotransmitter release and mediating neuronal excitability and plasticity given its pre- and postsynaptic localisation (Kaczmarek & Zhang, 2017; Rowan et al., 2016). Dysregulation of K_v_3.4 channel expression has been implicated in several neurological conditions due to it’s role in pathways linked to oxidative stress and hypoxia (Kääb et al., 2005; Song et al., 2017). K_v_3 channels are also involved in signalling pathways linked to neurodegenerative diseases such as spinal cerebellar ataxia (Zhang & Kaczmarek, 2016). In early and late stages of AD, K_v_3.4 gene expression was increased in a study of frontal cortex from 7 donors (2 control, 3 early AD and 2 late AD cases) (Angulo et al., 2004). In this same study, protein level measured by western blot was also increased in AD compared to control frontal cortex, and Kv3.4 staining with immunohistochemistry was observed in a punctate pattern in all cases, with some accumulation around plaques in AD samples (Angulo et al., 2004). Further, increased K_v_3.4 protein levels were observed in 3 Tg2576 transgenic mice (which develop amyloid plaques) compared to 3 wild-type mice (Angulo et al., 2004). A subsequent study in the same mouse line specifically demonstrated astrocytic upregulation of K_v_3.4 and that lowering K_v_3.4 levels caused downregulation of a reactive astrocyte marker and Aβ oligomers (Boscia et al., 2017). This observation may be linked with the notion that the K_v_3.4-mediated K^+^ efflux is implicated in the activation of astrocytic inflammasomes and reduced astrocytic phagocytosis in the early stages of AD (Boscia et al, 2017; Piccialli et al, 2020). High concentrations of Aβ (5 μM) applied to rat primary hippocampal neuronal cultures or differentiated PC12 cells also induced K_v_3.4 expression and morphological abnormalities which could be prevented by inhibiting K_v_3.4 channel activity (Ciccone et al., 2019; Pannaccione et al., 2007).

Here we tested the hypothesis that K_v_3.4 channels play an important role in pathways that mediate toxicity towards synapses in the Alzheimer’s brain and that reducing levels of this channel can protect synapses from Aβ-induced degeneration. To test this hypothesis, we examined dendritic spines in plaque-bearing APP/PS1 mice and littermate controls with and without lowering K_v_3.4 expression. Further we examine human iPSC-derived cortical neurons and human post-mortem brain tissue to determine whether K_v_3.4 is indeed expressed in synapses where it may mediate Aβ toxicity.

## Materials and Methods

### Mice

APPswe/PS1dE9 (APP/PS1, n=10) mice originally purchased from Jackson Laboratory were bred and aged in house. These double transgenic mice overexpress a human mutant amyloid precursor protein gene with the Swedish mutation and a human mutant Presenilin 1 gene with the deletion of exon 9 (Jankowsky et al, 2004). Wild-type (WT, n=6) littermates were used as controls. Both sexes of mice were used. All mice were group housed in a 12-hour day/night cycle with ad libitum access to food and water. All mice underwent surgery for stereotaxic injections at 7 months of age and were sacrificed 7 weeks post-injection for brain collections. Table 1 shows a summary of the mice used in this study. Experiments were performed in accordance with the UK Animal (Scientific Procedures) Act 1986 and the Directive 2010/63EU of the European Parliament and the Council on the protection of animals used for scientific purposes.

**Table 1.** Summary of mice used in this study. Only a subset of APP/PS1 mice were analysed for K_v_3.4 staining intensity. N.A.: Not applicable.

| Mouse ID | Genotype | Sex | Spine density<br>*Mean (N dendritic segments)* |  |  |  | K<sub>v</sub>3.4 intensity<br>(normalised to background)<br>*Mean (N cell bodies)* |  |
| --- | --- | --- | --- | --- | --- | --- | --- | --- |
|  |  |  | tdTomato (Control) |  | eYFP (K<sub>v</sub>3.4 knockdown) |  | tdTomato (Control) | eYFP (K<sub>v</sub>3.4 knockdown) |
|  |  |  | Far | Near | Far | Near |  |  |
| TS23 | WT | F | 1.82 (20) | N.A. | 2.38 (20) | N.A. | 3.10 (14) | 2.34 (10) |
| TS25 | WT | F | 2.11 (20) | N.A. | 2.52 (20) | N.A. | 2.65 (10) | 2.32 (10) |
| TS511 | WT | M | 1.59 (20) | N.A. | 2.37 (20) | N.A. | 2.79 (13) | 2.34 (16) |
| TS517 | WT | F | 1.77 (20) | N.A. | 2.42 (20) | N.A. | 2.30 (9) | 2.30 (11) |
| TS531 | WT | M | 2.07 (20) | N.A. | 2.12 (20) | N.A. | 2.46 (10) | 2.00 (14) |
| TS532 | WT | M | 0.80 (20) | N.A. | 1.17 (20) | N.A. | 1.85 (11) | 1.48 (10) |
| TS476 | APP/PS1 | F | 0.66 (20) | 0.58 (24) | 0.64 (20) | 0.48 (32) | - | - |
| TS478 | APP/PS1 | F | 0.72 (10) | 0.53 (37) | 0.66 (19) | 0.52 (30) | - | - |
| TS481 | APP/PS1 | M | 0.80 (12) | 0.61 (26) | 1.33 (23) | 0.79 (24) | - | - |
| TS499 | APP/PS1 | F | 0.48 (26) | 0.46 (37) | 0.49 (12) | 0.46 (29) | - | - |
| TS518 | APP/PS1 | F | 0.44 (13) | 0.48 (10) | 0.68 (39) | 0.85 (38) | - | - |
| TS520 | APP/PS1 | M | 1.43 (20) | 0.73 (20) | 1.20 (20) | 0.88 (20) | 2.40 (10) | 1.98 (14) |
| TS526 | APP/PS1 | F | 0.83 (20) | 0.50 (20) | 1.59 (20) | 1.17 (20) | 3.20 (10) | 2.43 (11) |
| TS533 | APP/PS1 | F | 2.04 (20) | 1.33 (20) | 2.26 (18) | 1.92 (23) | 3.22 (10) | 2.36 (11) |
| TS538 | APP/PS1 | M | 1.39 (20) | 0.57 (20) | 1.78 (20) | 1.37 (20) | 2.36 (11) | 2.17 (11) |
| TS546 | APP/PS1 | M | 1.45 (20) | 0.75 (18) | 1.72 (20) | 1.06 (20) | 4.81 (12) | 2.04 (12) |

### Viral Vectors

Adeno-associated viruses (AAVs) were used encoding the red fluorescent protein tdTomato, enhanced yellow fluorescent protein (eYFP) or the single guide RNA (sgRNA) that cleaves *K_V_3.4/KCNC4:* pENN.AAV.CAG.tdTomato.WPRE.SV40 (Addgene #105554), pAAV.CamKII(1.3).eYFP.WPRE.hGH (Addgene #10522), U6.sgRNA(mK_v_3.4).CMV.saCas9, respectively. Prior to use, 2 μL of pENN.AAV.CAG.tdTomato.WPRE.SV40 (1 × 10^13^ GC /mL) was diluted with an equal volume of 0.1M phosphate buffer saline (PBS) while 2 μL of pAAV.CamKII(1.3).eYFP.WPRE.hGH (1 × 10^13^ GC /mL) was mixed with 2 μL of U6.sgRNA(mK_v_3.4).CMV.saCas9 (1 × 10^11^ GC/mL).

### Stereotaxic surgery

At 7 months of age, mice were anaesthetised with isofluorane (3% for induction, 0.5-2% for maintenance). Fur on the head of the mice was shaved and Viscotears liquid gel was applied over both eyes before the animals were secured using ear bars in a stereotaxic apparatus. Body temperature was regulated using a heating pad and a rectal probe thermometer. After sterilising the surgical site with betadine and isopropyl alcohol, and performing local anaesthesia with subcutaneous injection of xylocaine (2 μg/g body weight), a 2-3 mm incision was made in the scalp to expose the skull. Burr holes were drilled in the skull, 1.5 mm bilaterally and 1 mm posterior to Bregma. Using a 10 μL Hamilton syringe, 4 μL of viral preparation was injected 0.7 mm deep into each burr hole on both somatosensory cortices at a rate of 420 nL/sec. Each hemisphere was randomly assigned to either experimental (K_v_3.4-knockdown) or control condition and received corresponding AAV viral injections into the somatosensory cortex (K_v_3.4-knockdown hemisphere: pAAV.CamKII(1.3).eYFP.WPRE.hGH: U6.sgRNA(mK_v_3.4).CMV.saCas9, 1:1; Control hemisphere: pENN.AAV.CAG.tdTomato.WPRE.SV40: PBS, 1:1). After injections, the scalp was sutured and mice were allowed to recover from anaesthesia in a heated chamber. Injected mice were singly housed under standard conditions until brains were harvested.

### Mouse brain tissue processing

After a 7-week incubation period to allow viral expression in neurons, the mice were euthanised and perfused transcardially with PBS followed by 4%paraformaldehyde (PFA; Agar Scientific, AGR1026). The brains extracted from the skull and post-fixed at 4° C for two hours before they were sectioned on a vibratome. Coronal sections of 50 μm thickness were collected between Bregma 0 and −2 to include the injection sites within the somatosensory cortices. Brain slices with eYFP (visualisation of K_v_3.4-knockdown condition) and tdTomato (control condition) expression were isolated after a quick examination under an epifluorescence microscope. These eYFP- and tdTomato-positive slices were then post-fixed in 4% PFA for another 20 minutes and stored in PBS at 4°C until use.

### Immunohistochemistry and Microscopy

For measurement of K_v_3.4 expression within injection sites, 50 μm floating slices were stained with an antibody specific to K_v_3.4 (Alomone Labs, APC-019) and an anti-rabbit Alexa Fluor Plus 647-conjugated secondary antibody (Invitrogen, A32795). After blocking the brain slices in 0.5% Triton-X 100 in PBS containing 3% normal donkey serum for an hour, samples were incubated with the primary antibody solution (1:200 dilution in blocking buffer) at 4°C overnight. The slices were washed in PBS and left in secondary antibody solution (1:5,000 dilution in blocking buffer) for two hours. To stain plaques, brain slices were mounted on microscope slides which were then dipped in 0.05% Thioflavin S in 50% ethanol for 8 minutes, followed by differentiation with 80% ethanol for 15 seconds. Immu-mount (Thermo Scientific #9990402) was applied before the samples were covered with a glass coverslip.

### Confocal imaging and analysis of mouse brains

To confirm K_v_3.4 reduction in the experimental versus control hemisphere, images stacks (61.5 μm × 61.5 μm × 10-28 μm with a z-step 0.3 μm, 63x zoom 3, N.A. 1.4) of tdTomato- and eYFP-positive cell bodies (n= ~10 images per mouse) were acquired on a Leica TCS SP8 confocal microscope using laser excitation at 488 nm for eYFP, 552 nm for tdTomato and 638 nm for K_v_3.4 staining with Alexa Fluor Plus 647 conjugated secondary. Using ImageJ (Schneider et al., 2012), the mean intensity of K_v_3.4 staining of cell bodies was measured. The K_v_3.4 staining intensities of cell bodies were normalised to background intensity measured in the same stack and z-section. Table 1 summarises the number of cell bodies analysed and the mean staining intensity of each animal.

To examine the effect of K_v_3.4-knockdown on dendritic spine densities, image stacks (61.5 μm x 61.5 μm x 3-25 μm with a z-step 0.3 μm, 63x zoom 3, N.A. 1.4) of dendritic segments, either labelled with eYFP or tdTomato, from cortical layer II, III and V pyramidal neurons were acquired. Only dendritic segments more than 20 μm in length were selected. For every APP/PS1 mouse, dendritic segments were captured for each condition: eYFP-positive, near a plaque (0-30 μm, using laser excitation 405 nm for ThioS staining); eYFP-positive, far from plaques (>30 μm); tdTomato-positive, near a plaque and tdTomato-positive, far from plaques. As WT mice do not present plaques, dendrite selection was made without regard to plaque proximity. Prior to image analysis, all images were randomly code-blinded by an observer and the quantification of spine densities was performed on greyscale images, thereby avoiding experimenter bias. Z-stack reconstructions and measurements were performed in ImageJ. The distance of dendrite segments to the nearest plaque, if present, was measured. The selected dendritic segments were traced in detail to count the number of spines present and their morphological class recorded, either as mushroom, thin, stubby or branched. Branched spines had more than one head (Harris et al, 1992; Peters & Kaiserman-Abramof, 1970; Spires et al, 2005b). Mushroom spines had a clear protrusion with head diameter ≥ 2x neck diameter. Thin spines had similar head and neck diameter (head diameter < 2x neck diameter), while stubby spines had no visible neck (head diameter < neck diameter). The length of dendrite segments were measured to calculate linear spine density (n spine per μm along the dendrite). Dendritic shaft diameters were also measured at each end and the midpoint of each segment to produce an average diameter. The number of dendritic segments analysed and the mean spine density of each animal are shown in Table 1.

### Human subjects

Use of human tissue for post-mortem studies has been reviewed and approved by the Edinburgh Brain Bank ethics committee and the ACCORD medical research ethics committee, AMREC (ACCORD is the Academic and Clinical Central Office for Research and Development, a joint office of the University of Edinburgh and NHS Lothian, approval number 15-HV-016). The Edinburgh Brain Bank is a Medical Research Council funded facility with research ethics committee (REC) approval (11/ES/0022). Use of human stem cell derived neurons was approved by the NHS Lothian REC (10/S1103/10).

### Western blots of human brain tissue

The concentrations of protein samples from human brain whole homogenate preparations were measured via the bicinchoninic acid (BCA) assay (Micro BCA protein assay kit, ThermoFisher Scientific). 20 μg of total protein were mixed 1:1 with Laemmli buffer and boiled at 95 °C for 10 minutes, and loaded to NuPAGE^™^ 4 to 12%, Bis-Tris gels. Gels were run at 120 V, 400 mA, >2 hours, until the loading dye ran to the bottom of the gel. Protein molecular ladder (Color Prestained Protein Standard, Broad Range 11–245 kDa, NEB, P7712S). Dry transfer was performed using an iBlot^™^ 2 gel transfer device (Invitrogen) (8.5 minutes, 20 V) onto a polyvinylidene difluoride (PVDF) membrane. Membranes were briefly washed in phosphate buffered saline (PBS) and then immersed in Revert^™^ 700 Total Protein Stain for 5 minutes at room temperature. Membranes were immediately washed in Revert 700 Wash Solution, exposed at 700 nm for 1 minute and imaged (Odyssey^®^ Fc, LICOR). Membranes were washed (Revert Destaining Solution) and blocked for 1 hour at room temperature (LICOR Blocking buffer mixed 1:1 with PBS enriched with 0.1% Tween-20 in PBS-T). Membranes were incubated with primary antibody (anti-KCNC4; Alomone #APC-019; 1:200) overnight at 4 °C in blocking buffer. Following 3x 10-minute washes in PBS-T, membranes were incubated with horseradish peroxidase-conjugated (HRP) secondary antibodies (goat anti-rabbit-HRP; 1:5,000) in blocking buffer for 1 hour, at room temperature. Membranes were washed 3x 30 minutes in PBS-T followed by incubation with Amersham^™^ ECL^™^ Prime Western Blotting Detection Reagent (Cytiva) to visualise bands. Chemiluminescent bands were visualised using the LICOR Odyssey^®^ Fc system. Tissue from nine donors was used for this study and their details are found in supplementary information.

**Supplemental table 1:** Human brain samples used for western blot analysis

|  | BBN | Sex | Age | PMI (hours) | Braak Stage |
| --- | --- | --- | --- | --- | --- |
| Braak 0 - 1 | 001.35138 | F | 73 | 74 | 0 |
|  | 001.34150 | M | 63 | 115 | 0 |
|  | 001.35420 | F | 82 | 114 | I |
|  | 001.35215 | M | 82 | 40 | I |
|  | 001.31504 | F | 65 | 76 | I |
|  | 001.31503 | F | 57 | 73 | I |
|  | 001.28402 | M | 79 | 49 | I |
|  | 001.28794 | F | 79 | 61 | I |
|  | 001.30208 | M | 73 | 66 | I |
|  | 001.30178 | M | 72 | 60 | 0 |
| Braak 3 - 4 | 001.32820 | F | 69 | 94 | IV |
|  | 001.29528 | F | 93 | 31 | III |
|  | 001.28411 | F | 88 | 65 | III |
|  | 001.28405 | M | 84 | 83 | IV |
|  | 001.26499 | F | 81 | 59 | III |
|  | 001.26491 | M | 77 | 27 | III |
|  | BBN_26492 | F | 74 | 71 | IV |
|  | BBN_24323 | M | 67 | 103 | IV |
|  | BBN_20994 | M | 75 | 59 | III |
|  | BBN_19600 | F | 85 | 36 | III |
| Braak 5 - 6 | 001.35183 | M | 74 | 75 | VI |
|  | 001.35096 | M | 72 | 103 | VI |
|  | 001.33698 | F | 90 | 76 | VI |
|  | 001.33636 | M | 93 | 43 | VI |
|  | 001.32929 | F | 85 | 80 | VI |
|  | 001.30883 | F | 61 | 69 | VI |
|  | 001.29911 | M | 66 | 52 | VI |
|  | 001.29521 | M | 95 | 96 | VI |
|  | 001.26500 | M | 81 | 83 | VI |
|  | BBN_24668 | F | 96 | 61 | VI |

#### Human iPSC-derived cortical neuron culture

We differentiated five iPSC lines from blood samples of healthy aged people participating in the Lothian Birth Cohort 1936 (LBC1936) study (Toombs et al., 2020). Briefly, peripheral blood mononuclear cells (PBMCs) were reprogrammed using non-integrating oriP/EBNA1 backbone plasmids expressing six iPSC reprogramming factors (OCT3/4 (POU5F1), SOX2, KLF4, L-Myc, shp53, Lin28, SV40LT). All lines demonstrated STR matched karyotype. Pluripotency was validated by immunocytochemistry, alkaline phosphatase staining, and Pluritest. Tri-lineage differentiation potential was confirmed by hPSC Scorecard and embryoid body formation techniques. All lines were confirmed to be mycoplasma negative. Tissue culture is conducted at 37°C, 5% CO_2_. iPSCs are maintained in 6 well culture plates coated with 1:100 Geltrex and fed daily with Essential 8 (E8) media. iPSCs were passaged with 0.1% EDTA pooled at a ratio of 5:1 in 6 well culture plates coated with 1:100 Geltrex and fed with E8 media. Differentiation into glutamatergic neurons was conducted by dual SMAD inhibition as described in (Supplemental figure 1 and (Shi et al., 2012)).

Generation of soluble human brain fraction was conducted according to a published protocol (W. Hong et al., 2018). Brain tissue was chopped and incubated for 30 min in artificial CSF (pH 7.4) supplemented with 1x complete mini EDTA-free protease inhibitor cocktail tablet (Roche, 11836170001). The was centrifuged at 2000 RCF for 10 mins to remove large, insoluble debris then centrifuged at 200,000 RCF for 110 mins. The resulting supernatant was dialyzed to remove salts and any drugs taken by the donor. The resulting homogenate was immunodepleted to remove soluble amyloid beta or mock immunodepleted using protein A agarose beads (Thermo, 20334) with 4G8 antibody (Biolegend, 800703) or mouse serum (Merck, M5905) as a mock condition (W. Hong et al., 2018). Concentration of Aβ1-42 in Aβ+ and Aβ-homogenate was quantified by sandwich ELISA (WAKO, 296-64401), according to manufacturer instructions. For experiments, Aβ+ homogenate was used at a final concentration of 15pmol/L. Aβ-homogenate approximated the ELISA lower limit of quantification, therefore Aβ content could not be accurately determined. Aβ-homogenate was diluted by the same factor as Aβ+, with a final concentration considered to be <1pmol/L.

iPSC-derived cortical neurons were fixed with 4% formalin (Polysciences, cat.04018-1) for 15 mins, rinsed in Dulbecco’s phosphate buffered saline (D-PBS, Thermo 14190250) and stored in D-PBS at 4°C for up to a month before staining. To stain, coverslips containing fixed neurons were permeabilized and non-specific antigens blocked by incubating in D-PBS with 0.2% Triton-X (Merck, cat. X100-500ML) and 10% donkey serum (Merck, cat.566380-10ML) for one hour. Coverslips were incubated overnight at 4°C with primary antibodies: Homer (rabbit anti-Homer; Abcam; 1:500), MAP2 (guinea pig anti-MAP2; Synaptic Systems 188004; 1:1000), Kv3.4A (sheep anti-Kv3.4A; Autifony Ab101; 1:100) diluted in D-PBS containing 0.2% Triton-X and 1% donkey serum. Cells were then washed with D-PBS containing 0.1% Triton-X, and incubated in secondary antibodies diluted 1:500 in D-PBS containing 0.1% Triton-X and 1% donkey serum for one hour (donkey anti-rabbit-488 (Abcam ab150073), donkey anti-guinea pig-594 (Merck, SAB4600096), donkey anti-sheep-647(Thermo A21448)). Coverslips were mounted on slides (VWR, 631-0847) with mounting media (Merck, cat.345789-20ML) and imaged on a Leica TCS confocal microscope with an oil immersion 63x objective. Ten fields per coverslip containing MAP2 staining were randomly selected for imaging and image stacks acquired through the thickness of the cell layer (0.3μm per step). Image stacks were processed using custom software to segment staining, calculate the density of synaptic markers along MAP2 positive processes, colocalization of K_v_3.4 with Homer 1 post-synaptic terminals, and measure the intensity of K_v_3.4 staining in dendrites. All image analysis scripts are freely available at https://github.com/Spires-Jones-Lab.

Aged neurons were incubated with Trizol (Thermo, 15596026) for five mins. Neurons were homogenised with a P200 pipette and collected in DNase/RNase free tubes (Eppendorf, 30108051). 200uL of chloroform (Sigma, 288306-100ML) was added per 1mL Trizol sample and mixed by inversion. Samples were centrifuged at 12,000 RCF for 15 minutes at 4°C. The aqueous phase containing RNA was collected and an equal volume of 100% isopropanol added. The sample was centrifuged at 12,000 RCF for 10 mins at 4°C and the supernatant discarded. The pellet was washed in two cycles of 0.5mL of 70% ethanol and centrifugation at 12,000 RCF for 10 mins at 4°C. The sample was air dried for 10 mins at room temperature to evaporate any residual ethanol. The RNA pellet was solubilised in 30uL DEPC water (Thermo, AM9906) and transferred to a fresh DNase/RNase free Eppendorf, and stored at −80°C. Concentration and purity were measured using an LVis plate on a ClarioSTAR Plus spectrophotometer (BMG Labtech).

RT-qPCR was conducted using one-step RT-qPCR kit (Promega, A6020), according to manufacturer instructions. This product uses advanced BRYT green dye, enabling an RNA detection range of 500fg and up to 100ng, and alleviates the need for prior cDNA translation. Briefly, 100ng RNA samples were added to a mastermix preparation (GoTaq qPCR Mastermix (2X), GoScript RT mix for 1-step RT-qPCR (50X), Forward Primer (200nM), Reverse Primer (200nM), and DEPC H2O) to a volume of 20μL per well. Primers are described in Supplementary table XX. RT-qPCr was conducted in a thermal cycler (BioRad, CFX96 Touch Real-Time PCR Detection System), with an initial denaturation step (95°C, 10 mins), then cyclic denaturation (95°C, 10 seconds), annealing (60°C, 30 seconds), and extension (72°C, 30 seconds) for 40 cycles. This was concluded with a melt curve 65-95°C. Data were analysed with BioRad CFX Maestro software. Target expression was normalised to that of two reference genes (GAPDH and RPLP1) using ΔΔC_q_ method. GAPDH and RPLP1 were determined to be the most stable reference genes of a gene panel tested on iPSC-neurons treated with Aβ+ and Aβ− homogenate, with a Normfinder stability score of 0.09.

### Data analysis

Linear mixed effects models were used to analyse data with mouse as a random effect to account for multiple measurements per mouse. Genotype, treatment (K_v_3.4 knockdown or control), sex, and plaque proximity (in APP/PS1 mice only) were fixed effects in mouse analyses. Assumptions of the model fit were tested by visual inspection of residual plots. Where needed, the data were transformed to better fit model assumptions (transformations noted in results). Analysis of variance tests were run on linear mixed effects models to examine main effects and estimated marginal means with Tukey corrections were used for post-hoc group comparisons. For spine morphology data, Pearson’s chi-squared tests were used to compare all spines in each treatment group. For correlations with. Plaque distance, repeated measures correlations were conducted for each treatment group separately using a published package (Bakdash & Marusich, 2017). All statistical analyses were run in R Studio and analysis scripts, data files analysed, and analysis outputs including residual plots and Q-Q. plots for models are provided as supplemental information. Data are presented as box plots of data from all images analysed with individual data points showing a mean for each mouse or human to show biological variability.

## Results

To test the hypothesis that K_v_3.4 downregulation ameliorates dendritic spine loss associated with amyloid pathology, APP/PS1 mice and wildtype control mice were injected with an AAV containing the sgRNA targeting *K_v_3.4/KCNC4* into neurons of their somatosensory cortex, thereby decreasing K_v_3.4 levels in this region. This AAV was co-injected with an AAV that introduced the gene for eYFP which allowed fluorescent visualisation of neurons. The contralateral hemisphere which was injected with an AAV to express the tdTomato reporter in cells served as a within-animal control condition. A total of 10 APP/PS1 mice and 6 wildtype mice were used in this study. The K_v_3.4-knockdown by AAV was assessed by measuring fluorescence intensity of K_v_3.4 staining in a subset of the animals used (n=5 APP/PS1; n=6 WT mice). To examine synapse loss, dendrites labelled with eYFP/tdTomato were investigated in all mice, near (0-30 μm) and far (>30 μm) from plaques (if present), using confocal microscopy and measurements for dendritic spine density were produced using image stacks.

We observed a 21% decrease in K_v_3.4 staining intensity in eYFP-expressing neurons in which K_v_3.4 was knocked down compared to tdTomato-expressing control neurons (Figure 1, linear mixed effects model on Tukey transformed data followed by ANOVA, F[1,39.37]=38.22, p<0.0001). There was no significant effect of sex or genotype, nor was there an interaction between treatment and genotype.

**Figure 1.**
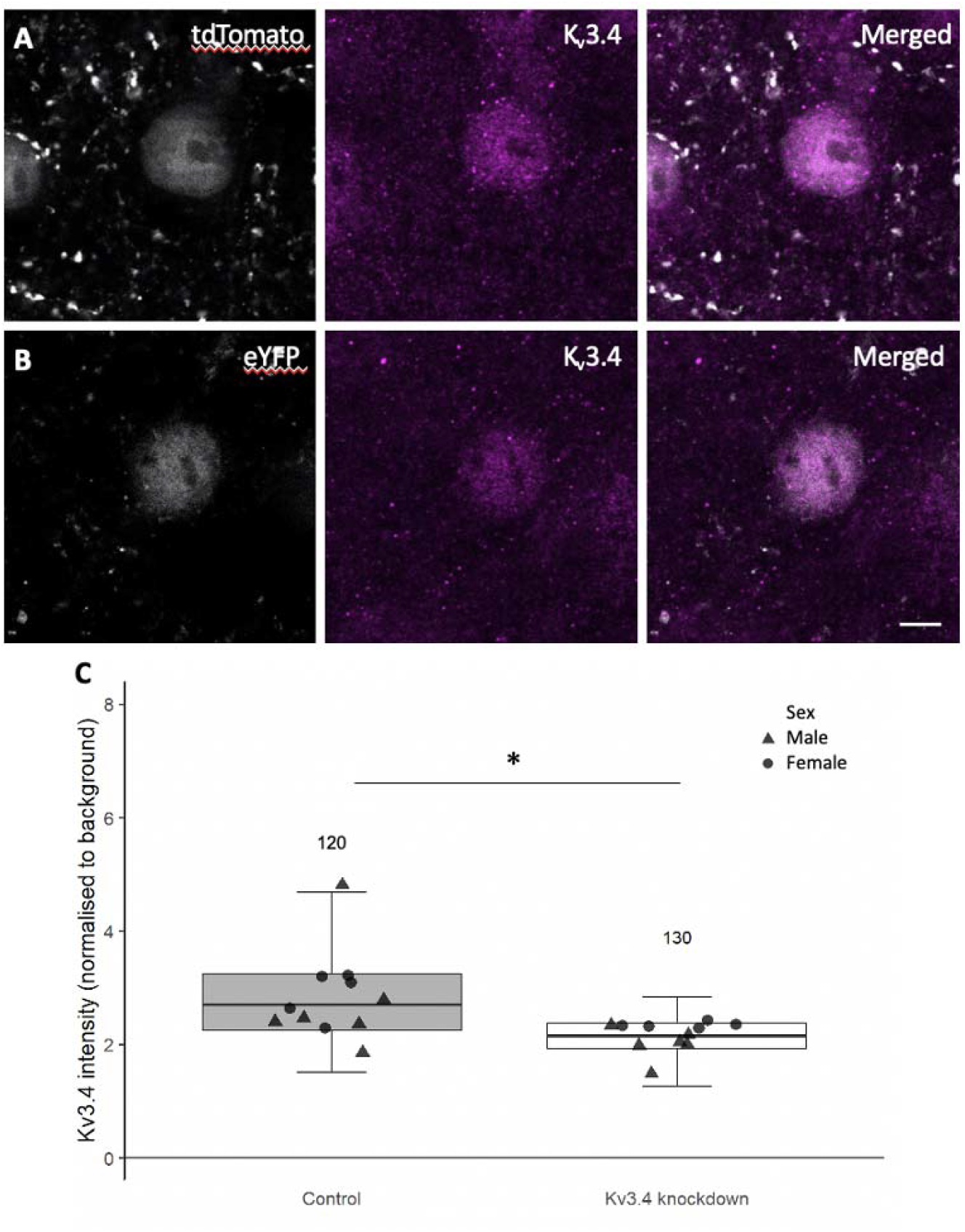
Representative immunostaining images of K_v_3.4 in neurons expressing either tdTomato or eYFP (A & B). For comparison purposes, tdTomato and eYFP cell bodies are visualised in grey pseudocolour and Kv3.4 staining in magenta. Compared to the tdTomato-filled neuron, the eYFP-filled neuron shows lower K_v_3.4 immunoreactivity. Analysis of K_v_3.4 immunostaining intensity revealed a 21% reduction in staining intensity in eYFP-filled cells relative to tdTomato-filled cells (C), confirming the action of U6.sgRNA(mK_v_3.4).CMV.saCas9 at knocking down K_v_3.4 levels. * p<0.0001 ANOVA effect of treatment; N above error bars represent number of cells analysed for each condition. Individual data point shows mean per mouse. Scale bar: 5 μm.

A reduction of dendritic spine density near plaques has been consistently reported in AD mouse models, including APP/PS1 mice used in our current study (Koffie et al., 2009; Moolman et al., 2004; Rozkalne et al., 2011). Here, to analyse the effects of K_v_3.4 reduction on plaque associated spine loss, we measured dendritic spine density of dendrite branches from layer II, III and V originating from cortical pyramidal neurons filled with eYFP/tdTomato in APP/PS1 and control mice (Figure 2A-F). In general, the spine density in APP/PS1 transgenic mice was significantly lower compared to wildtype mice (Figure 3A, F[1,13.06]=16.20, p=0.001). Under control conditions, dendrites in APP/PS1 mice had significantly less spines than those in wildtype mice (β=0.85, t=3.52, p=0.012, post-hoc estimated marginal means comparison), which is in accord with previous studies (Koffie et al., 2009; Moolman et al., 2004; Rozkalne et al., 2011). There is further a positive correlation between plaque distance and spine density in the tdTomato filled control dendrites of APP/PS1 mice (Figure 3C, repeated measures correlation r_rm_=0.14, df=221, 95% CI 0.014-0.272, p=0.03), which was absent in their eYFP filled K_v_3.4-knockdown dendrites (Figure 3C, r=0.095, df=245, 95% CI −0.031-0.271, p=0.14).

**Figure 2.**
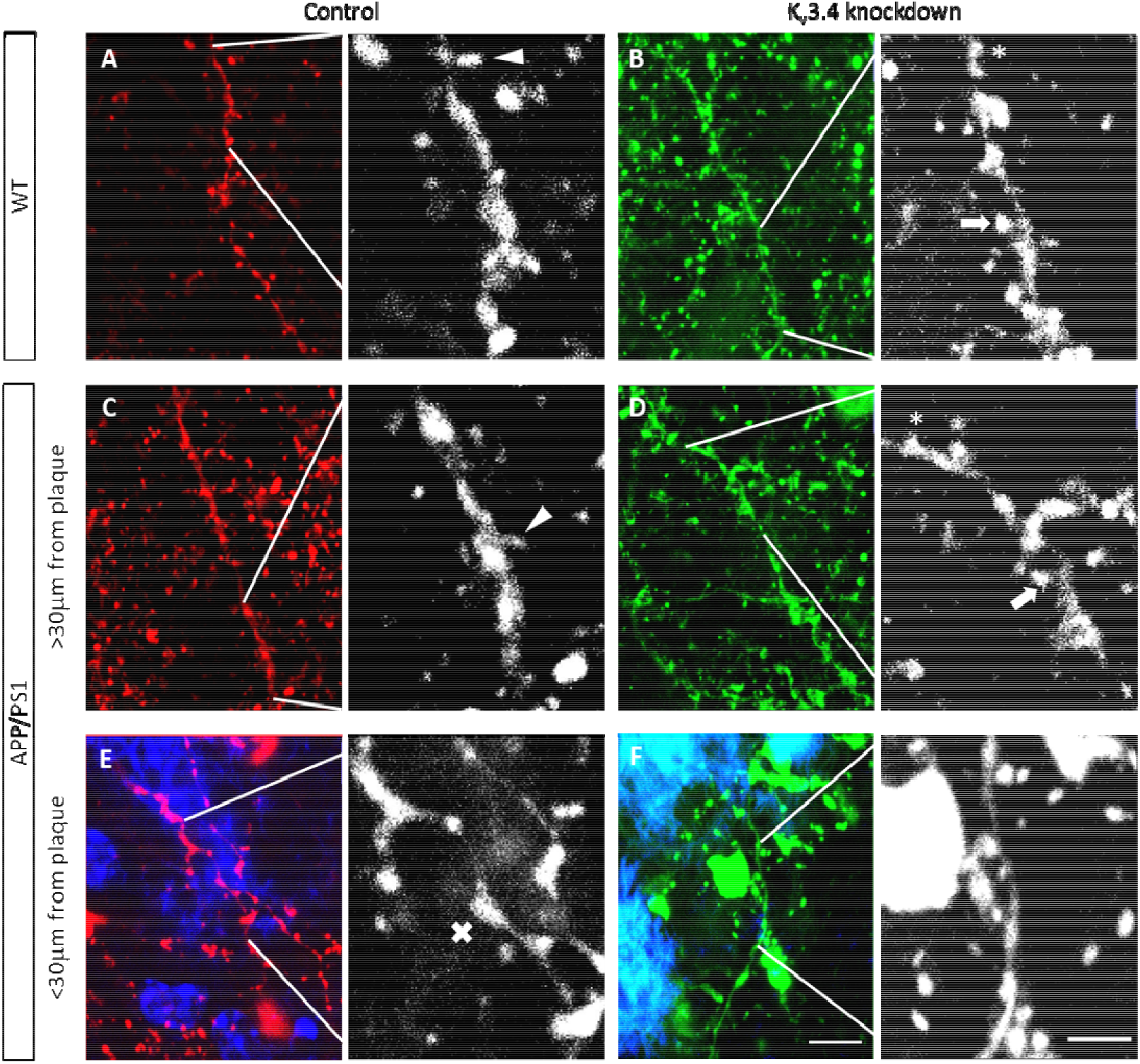
Dendritic spines were examined on dendrites from cortical pyramidal neurons filled with tdTomato or eYFP under control (A, C & E) or K_v_3.4 knockdown conditions (B, D & F), respectively. Spines were classified by shape as either mushroom (arrows), thin (arrowheads) or stubby (asterisks) based on the head to neck diameter ratio. Senile plaques in APP/PS1 mice were labelled in blue using ThioS (E &F). There is visible focal swelling (crosses) of dendritic segments that are in close proximity to plaques (E & F). Images are shown as maximum intensity Z-projections of 9 serial confocal images. Scale bar: 5 μm (left); 2 μm (right).

**Figure 3.**
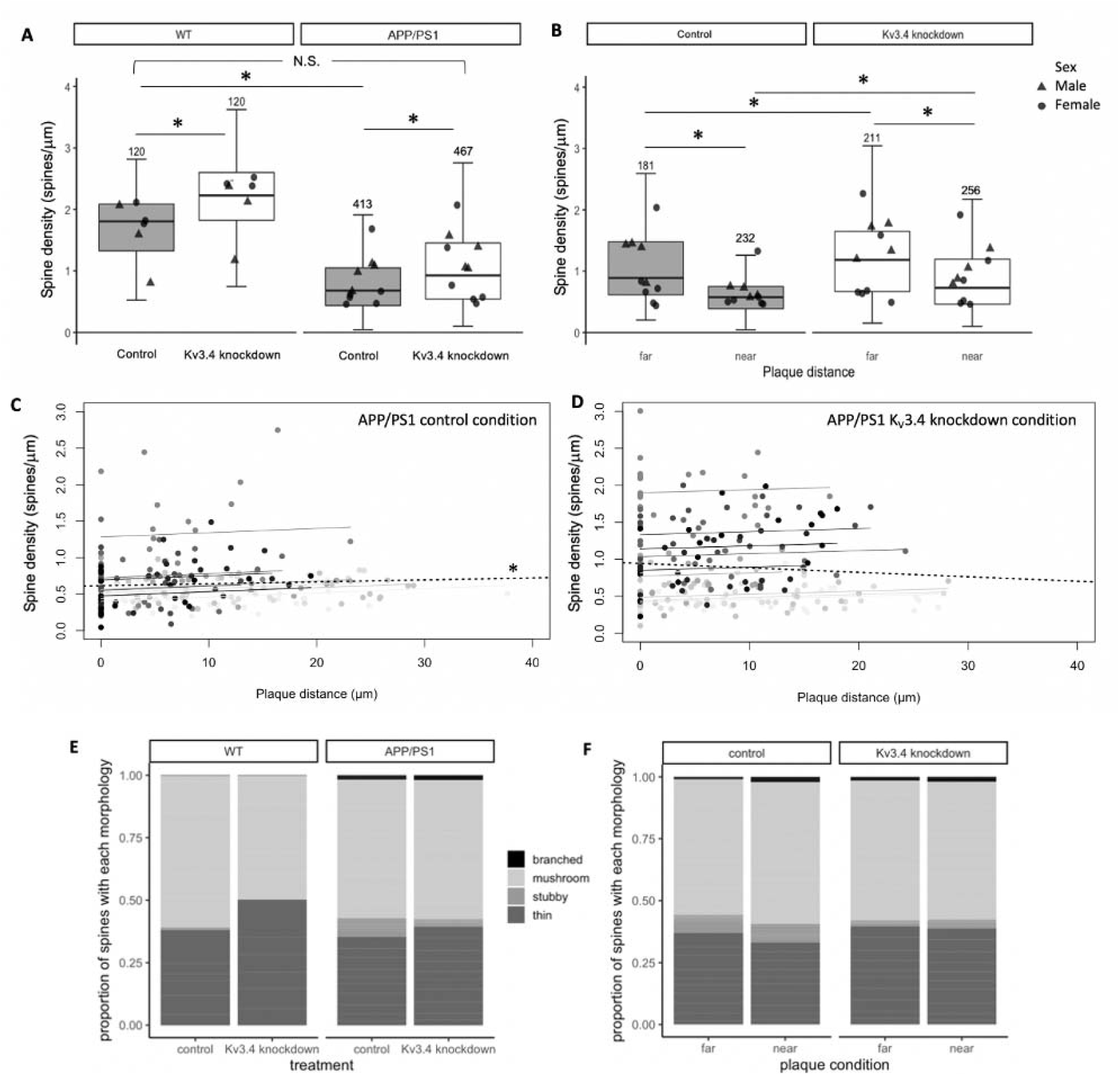
Under control conditions, there is a 62.4% reduction in spine density in APP/PS1 mice compared to WT mice without the APP and PSI transgenes (A). K_v_3.4 knockdown increases spine densities in both wildtype (WT) and APP/PS1 mice relative to control conditions, and restores spine densities in the transgenic mice to near WT control levels. In APP/PS1 mice, lower spine densities were recorded in dendrites within 30 μm of plaques in contrast to those distant from plaques (B). K_v_3.4 downregulation improved spine densities in all dendrites measured in APP/PS1 mice (A,B *p<0.05, pot-hoc estimated marginal means comparisons with Tukey correction. N above error bars represent number of dendrites analysed for each condition. Individual data point shows mean per mouse). Under control conditions, spine density of APP/PS1 mice correlated with distance from the nearest plaque (C, * p=0.03 repeated measures correlation). This correlation was absent in the K_v_3.4 knockdown dendrites, indicating that K_v_3.4 knockdown is protective (D). In C and D data points show individual dendrites shaded to show those from each individual mouse, the regression line for each mouse is shown in the same shade, and the overall regression is the black dotted line. The control hemisphere of APP/PS1 mice shows a shift in spine morphology to favour stubby spines (E, χ^2^= 18.15, p=0.03, Bonferroni adjusted post-hoc test), which was ameliorated by K_v_3.4 knockdown both compared to wild-type mice (E) and near and far from plaques (F). Values are reported as the proportion of spines in each category generated from means per mouse.

K_v_3.4 downregulation evidently increased spine density compared to the control hemisphere in both APP/PS1 and wildtype mice (Figure 3A, treatment effect F[1, 1102.12]= 163.29, p<0.0001) by 36.3% and 23.2%, respectively, with a bigger effect recorded in wildtype mice (Genotype x treatment interaction F[1,1102.12]=17.86, p<0.0001). Importantly, K_v_3.4 reduction in APP/PS1 mice restored spine density to be not significantly different from WT control level, as confirmed by post-hoc estimated marginal means comparisons (Figure 3A).

In APP/PS1 mice, dendrites within 30 μm of a plaque edge had lower spine densities than those farther away in both conditions (Figure 3B, effect of plaque proximity F[1,867.22] = 172.88, p<0.0001). In comparison to the control condition, K_v_3.4 knockdown significantly increased spine densities (effect of treatment, F[1,867.42] = 94.15, p<0.0001) by 32.7% in dendrites far from plaques (t= −5.78, p<0.001 post-hoc comparison) and by 26.4% in dendrites near plaques (t= −8.59, p<0.0001). This rescue of plaque-associated spine loss with K_v_3.4 knockdown is also illustrated when spine density is plotted versus plaque distance (Figure 3C).

In addition to spine densities, we also examined dendritic spine morphology, which affects postsynaptic integration of signals. There is a differential distribution of spine shapes in the control hemisphere of APP/PS1 mice with 7x more stubby spines compared to WT control level (Figure 3E, χ^2^ p<0.05 and χ^2^ Bonferroni adjusted post-hoc test p<0.05). K_v_3.4 knockdown reduces the number of stubby spines by more than half in these APP/PS1 mice (Figure 3E), which is also reflected in Figure 3F, where dendrites in both hemisphere conditions were further classified according to their plaque proximity. For dendrites distant from plaques and near plaques, the decrease in stubby spines in K_v_3.4 knockdown hemisphere relative to control hemisphere appears to be accompanied by a compensatory increase in the proportion of thin spines, but neither of these changes reached significance (Figure 4B).

**Figure 4.**
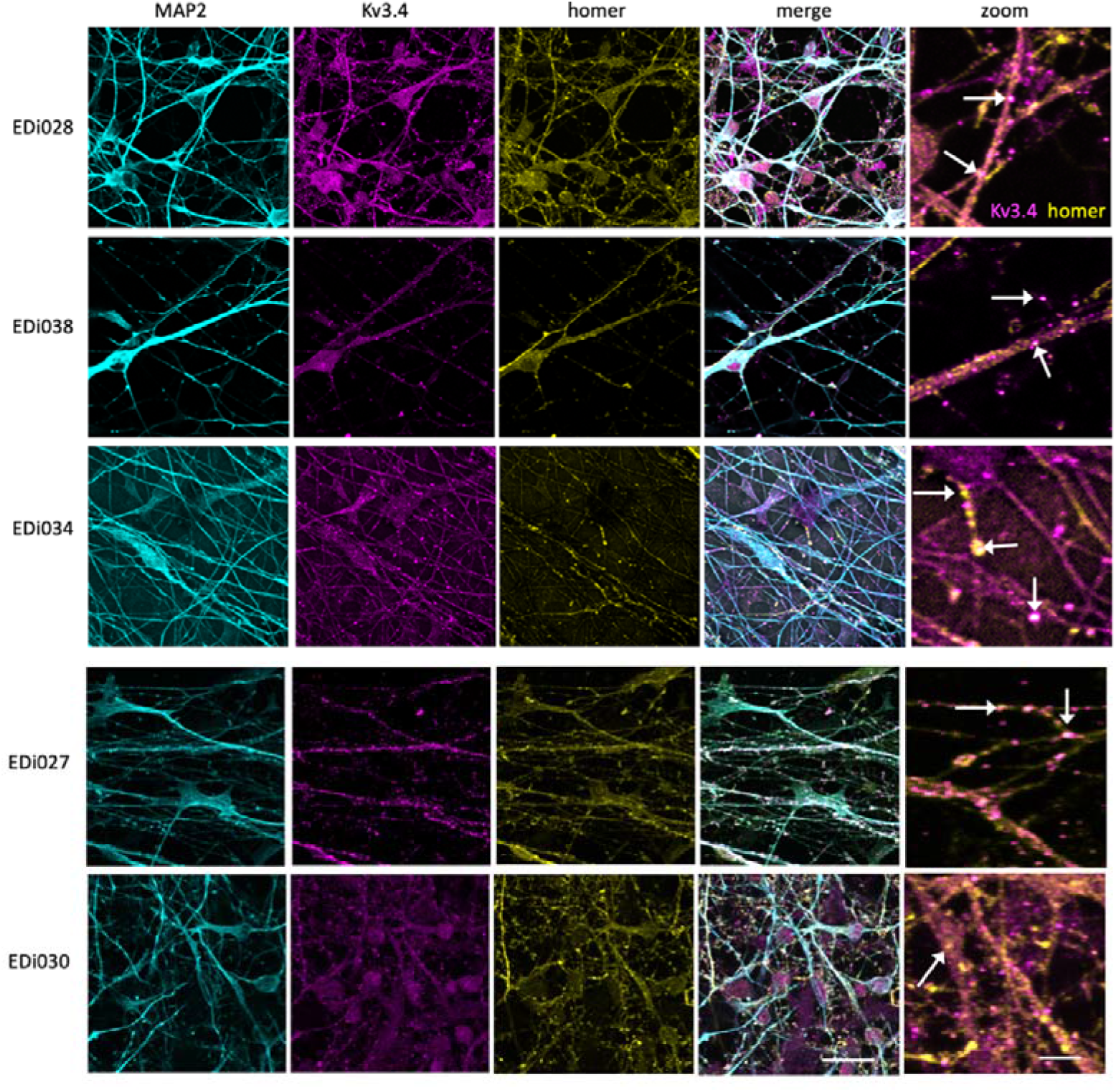
iPSC-derived cortical neurons from 5 human donors (lines EDi027, EDi028, EDi030, EDi034, EDi038) stained with MAP2 to label dendrites (cyan), postsynaptic protein homer 1 (yellow) and K_v_3.4 (magenta), have K_v_3.4 positive synaptic puncta along dendrites (arrows in zoom). Scale bar represents 20 μm (5 μm in zoom on right column).

To determine whether the rescue observed in our transgenic mouse line may be relevant to human brain, we examined K_v_3.4 in human post-mortem brain samples and human iPSC-derived neuronal cultures. In human iPSC neurons derived from blood samples from 5 different donors, we observe K_v_3.4 in synapses along dendrites (Figure 4), demonstrating the presence of K_v_3.4 protein in human synapses. Interestingly, challenging these neurons with human AD brain homogenate causes a decrease in K_v_3.4 expression alongside an increase in curvature of MAP2 positive neurites (Figure 5, ANOVA of linear mixed effects model of data transformed with the formula(Tortuosity-1) ^*1/7* to fit assumptions of model: F[1,6944]=15.21, p<0.0001). In human brain from people with very low (Braak 0-I), moderate (Braak III-IV) and extensive (Braak V-VI) Alzheimer’s disease pathology, we confirm that K_v_3.4 is expressed in both frontal and temporal cortices (Brodmann areas 9 and 20/21, respectively). In our samples, we do not observe any difference in levels between Braak stages or brain regions (linear model with Braak stage group, brain region, age, sex, and post-mortem interval as fixed effects: effect of Braak stage group F[2,50]=0.092, p=0.91; effect of brain region F[1,50]=0.626, p=0.43, data Tukey transformed to meet assumptions of linear model). There were also no effects of sex, age or PMI in our analyses (Supplemental Figure 1).

**Figure 5:**
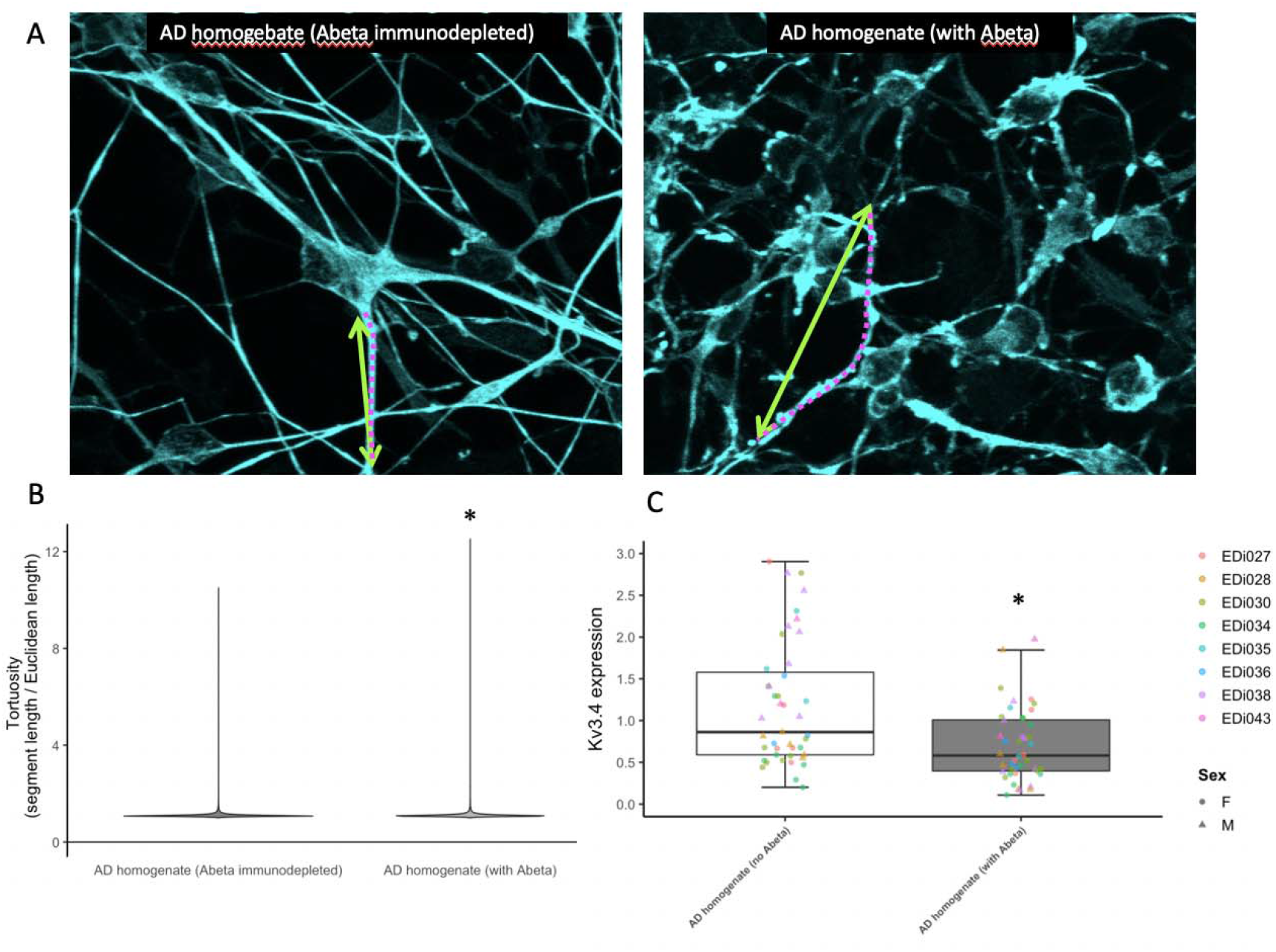
Human iPSC derived neurons were challenged with homogenate of brain from AD patients either mock immunodepleted for amyloid beta (Abeta) or immunodepleted to remove toxic Abeta. These neurons were fixed and stained for dendrites (MAP2, cyan) and dendrite tortuosity was measured as the dendritic segment length (magenta dotted line) divided by the direct Euclidean distance between the ends of the segment (green arrows, A). A violin plot of tortuosity shows an increase in curvature of dendrites treated with Aβ containing AD brain homogenate (B, * ANOVA on linear mixed effects model of transformed data with experiment and image nested in line as a random effect, F[1,6944]=14.21, p<0.0001). AD brain homogenate also causes a decrease in Kv3.4 expression as measured by qPCR (C, * ANOVA on linear mixed effects model with experiment nested in line as a random effect, F[1,47]=25.55, p<0.0001). Scale bars represent 20 μm in A and 5 μm in B.

**Supp Fig 1:**
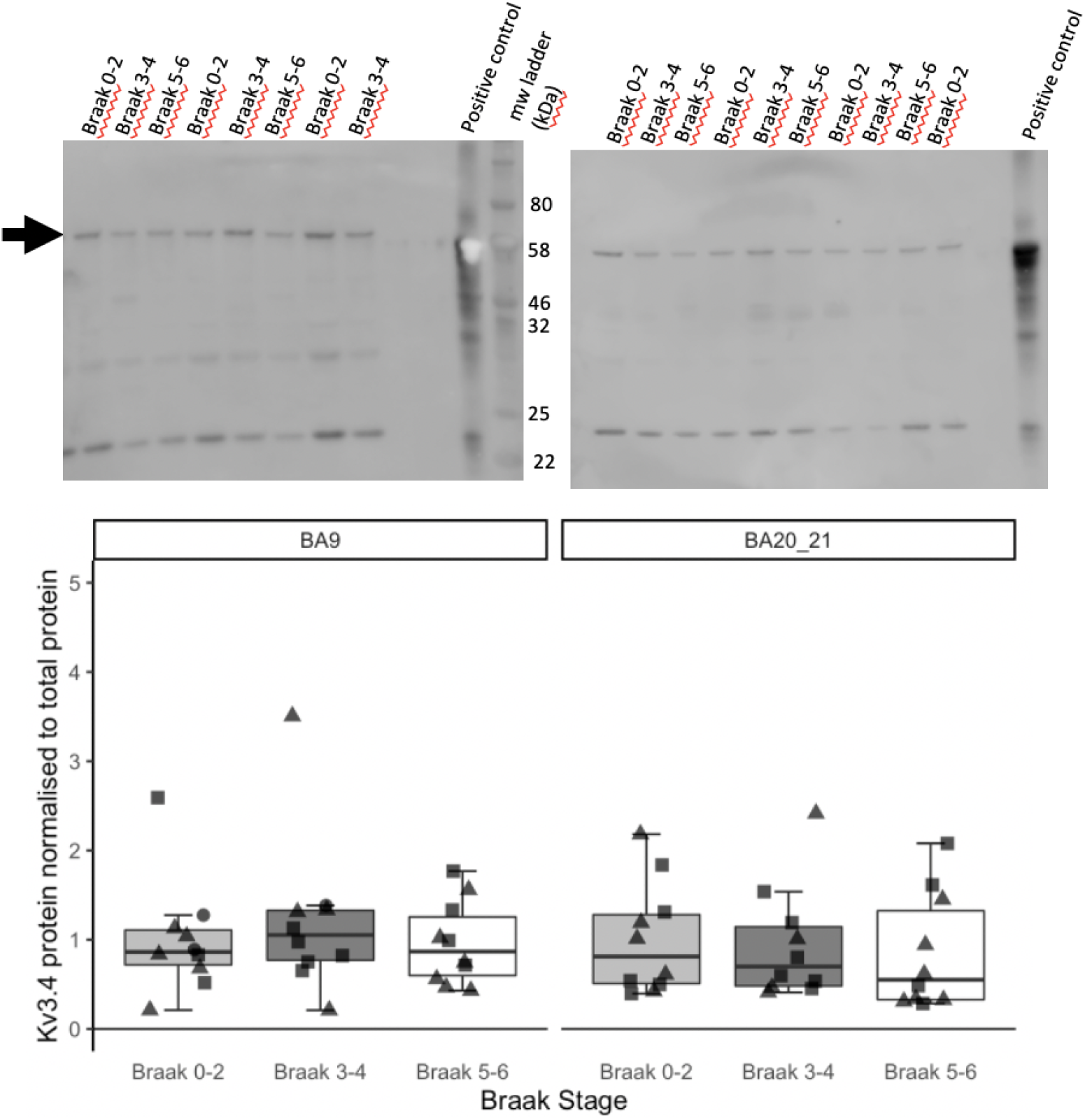
Western blot of human brain samples from frontal (BA9) and temporal (BA20/21) cortex of people with low (Braak 0-2), moderate (Braak 3-4) and extensive (Braak 5-6) Alzheimer’s disease pathology reveals no difference between brain regions or Braak stages (n=10 per group, linear model on Tukey transformed data p>0.05 for all fixed effects). The arrow indicates full-length K_v_3.4 protein immunoreactivity.

## Discussion

The loss of synapses has implications for the impaired learning and memory in Alzheimer’s patients considering the strong correlation between synapse loss and cognitive decline in disease (DeKosky et al., 1996; DeKosky & Scheff, 1990; Terry et al., 1991). Further the plasticity of synapses and ability to target synaptic receptors makes them attractive therapeutic targets. Based on previous data showing elevated expression of K_v_3.4 around Aβ plaques in human AD brain and in model systems (Angulo et al., 2004; Boscia et al., 2017; Pannaccione et al., 2007), here we tested the hypothesis that K_v_3.4 is involved in synapse loss in the APP/PS1 mouse model of amyloidopathy. Previous work using K_v_3.4 downregulation, either through siRNA or a selective toxin blocker, has shown neuroprotective effects in hippocampal cultures and in Tg2576 mice (Boscia et al, 2017; Ciccone et al, 2019; Pannaccione et al, 2007). In this study, we observed that the downregulation of K_v_3.4 using CRISPR/Cas9 AAV ameliorates plaque-associated dendritic spine loss in APP/PS1 mice. Upon examination of cortical pyramidal neurons, we found dendritic spine loss in APP/PS1 mice which is exacerbated near plaques, in agreement with previous findings in several different plaque-bearing transgenic mice (Koffie et al., 2009; Moolman et al., 2004; Rozkalne et al., 2011; Spires, 2005). K_v_3.4 knockdown rescues this phenotype in APP/PS1 mice, restoring spine density to wildtype control levels. Given their ability to impact membrane depolarisation and neurotransmitter release it certainly anticipated that the fine control of potassium (and indeed other) channel expression at synapses will act as a regulator of synaptic activity in several brain regions. Dendritic spines make up the postsynaptic element of over 90% of cortical excitatory synapses, thus our findings indicate that K_v_3.4 downregulation alleviates synapse loss in plaque-bearing mice. Although we did not replicate published findings (Angulo et al 2004) showing increased K_v_3.4 levels in human AD brain compared to controls, we do observe K_v_3.4 specifically localised to synapses in human iPSC-derived cortical neurons, supporting the possibility that K_v_3.4 in human synapses may mediate Aβ-induced synaptotoxicity.

One potential mechanism linking Aβ to K_v_3.4 is that oligomeric Aβ promotes the generation of Ca^2+^-induced reactive oxygen species (ROS) and activates the transcriptional factor nuclear factor kappa-B (NF-_K_B) that in turn upregulates K_v_3.4 gene expression. The upregulated expression of K_v_3.4 mediates excessive K^+^ efflux, leading to the activation of caspase-3 (Piccialli et al, 2020). Caspase-3 has been implicated in spine degeneration and consequent synaptic failure (D’Amelio et al., 2011) as well as to the accumulation of tau in neurofibrillary tangles (de Calignon et al., 2010; Spires-Jones et al., 2008). Caspase-3 activated calcineurin has been shown to drive the internalisation of α-Amino-3-hydroxy-5-methyl-4-isoxazolepropionic acid (AMPA) receptors from postsynaptic sites, and this is sufficient to cause spine elimination and the loss of synaptic NMDA receptors (D’Amelio et al., 2011; Hsieh et al., 2006; Snyder et al., 2005). Our previous work in mouse models has demonstrated that lowering calcineurin levels is also protective against dendritic spine loss in mouse models (Rozkalne et al., 2011; Wu et al., 2010). Elevated ROS production, NF-_K_B activity and K^+^ efflux are also prerequisites for the formation of microglial and astrocytic inflammasomes (Venegas & Heneka, Michael, 2019), and data are accumulating linking both microglia and astrocytes to synapse loss in AD models (Henstridge et al., 2019; S. Hong et al., 2016; Litvinchuk et al., 2018). It is also noteworthy that Kv3.4 is expressed in astrocytes in addition to neurones where it is upregulated in models of AD (Boscia et al 2017). Hence a direct impact of Kv3.4 downregulation on the reactive astrocytes in the AD brain may also deliver an additional and useful benefit alongside the impact on neuronal synaptic health quantified in detail here. Another potential explanation for the spine density recovery observed in our APP/PS1 mice is that reducing K_v_3.4 may have an effect on synapses independent of Aβ. This is supported by. Our data showing that knockdown of K_v_3.4 also increased spine density in WT mice.

In addition to the impact on synapse density, we also observed changes in the proportion of specific spine types, noting a clear increase in the number of stubby spines in APP/PS1 mice. This observation has been reported in numerous studies using cortical biopsies from AD patients, and transgenic mice carrying a familial-AD associated mutant *APP* transgene (Androuin et al., 2018; Spires-Jones et al., 2007; Tackenberg & Brandt, 2009). Studies in hippocampal cultures as well as *in vivo* in mice further revealed a gradual change from mushroom to stubby spines upon Aβ (Penazzi et al., 2016), highlighting the impact of Aβ on dendritic spine dynamics. While stubby spines are suggested to be the morphological correlate of LTD induction downstream of Aβ (Li et al., 2009), the loss of mushroom spines is seen as a morphological marker for synaptic failure (Tackenberg & Brandt, 2009). The increase in stubby spines without significant changes in mushroom spines observed our current study has also been documented in a recent hippocampal slice culture study with acute Aβ treatment (Ortiz-Sanz et al., 2020). In contrast to mature mushroom spines that form strong synaptic connections, stubby spines are immature, more dynamic and relatively scarce in the mature brain (Fiala et al., 1998; Harris et al., 1992). Spine outgrowth and maturation are dependent on NMDA and AMPA receptors (Engert & Bonhoeffer, 1999; Matus, 2000). Assuming that the spine loss detected in our APP/PS1 mice was a result of caspase-3 induced AMPA receptor endocytosis, the resulting loss of both AMPA and NMDA receptors may explain the increased number of immature stubby spines in our transgenic mice. Fundamentally, changes in spine morphology affect the postsynaptic integration of signals. While the volume of the spine head is important for the expressions of NMDA and AMPA receptors (Matsuzaki et al., 2001, 2004), the spine neck is responsible for Ca^2+^ compartmentalisation (Grunditz et al., 2008). Ca^2+^ imaging studies demonstrated a drastic increase in accumulated Ca^2+^ at the base of short stubby spines devoid of a neck (Noguchi et al., 2005). This would permit such a copious amount of Ca^2+^ to enter the parent dendrite that causes a loss of spine-to-dendrite Ca^2+^ homeostasis, as observed through Ca^2+^ imaging in APP/PS1 mice (Kuchibhotla et al., 2008). Our observation that K_v_3.4 knockdown shifts spine morphology from stubby to thin in APP/PS1 mice is interesting as thin spines become less prevalent alongside age-related cognitive deterioration in monkeys (Dumitriu et al., 2010). Thin spines are thought to be “learning spines” that are capable of strengthening plasticity in the local circuit (Bourne & Harris, 2007). Thus, the potential of targeting K_v_3.4 to increase the proportion of thin spines in AD may also result in restored plasticity and cognitive benefits, as reported in the 17β-estradiol treatment study that targets spine dynamics in rhesus monkeys (Hao et al., 2006).

Taken together, our results demonstrate that K_v_3.4 downregulation is able to reduce dendritic spine loss and restore spine density and morphology in aged APP/PS1 mice. We also observe K_v_3.4 expression on the synapses of human neurons making it a promising target for the development of novel therapeutic agents that seek to modulate Kv3.4 expression and/or function for the treatment of Alzheimer’s disease, and potentially other CNS diseases.

## Acknowledgements

This work was funded by Autifony Therapeutics Ltd, Alzheimer’s Research UK (ARUK-TVPG2018-010), the UK Dementia Research Institute, which receives its funding from DRI Ltd, funded by the UK Medical Research Council, Alzheimer’s Society, and Alzheimer’s Research UK, and the European Research Council (ERC) under the European Union’s Horizon 2020 research and innovation programme under grant agreement No 681181, and NIH grant R56-AG072473 (M.J.M.R.), and the Emory Alzheimer’s Disease Research Center Grant 00100569 (M.J.M.R.).

